# A mental number line in human newborns

**DOI:** 10.1101/159335

**Authors:** Rosa Rugani, Marco Lunghi, Elisa Di Giorgio, Lucia Regolin, Beatrice Dalla Barba, Giorgio Vallortigara, Francesca Simion

## Abstract

In the 19^th^ century Francis Galton first reported that humans represent numbers on a mental number line with smaller numbers on the left and larger numbers on the right. It has been suggested that this orientation emerges as a result of reading/writing habits for both words or numbers. Recent evidence in animals and infants in the first months of life has challenged the primary role of language in determining the left-to-right direction of spatial-numerical association, SNA. However, the possibility that SNA is learnt by early exposure to caregivers’ directional biases is still open. Here we show that 55-hour-old newborns, once habituated to a number (i.e., 12), spontaneously associated a smaller number (i.e., 4) with the left side and a larger number (i.e., 36) with the right side of space. Moreover, SNA in neonates was not absolute but relative. The same number (i.e., 12) was associated with the left side whenever the previously experienced number was larger (i.e., 36), but with the right side whenever the number was smaller (i.e., 4). Control on continuous physical variables showed that the effect was specific of discrete magnitudes. Hence, soon after birth humans associate smaller numbers with the left space and larger numbers with the right space. These results constitute strong evidence that in our species SNA originates from pre-linguistic and biologically precursors in the brain.

**SIGNIFICANCE STATEMENT:** For human adults, the representation of number and space is profoundly intertwined. Humans represent numbers on a left to right oriented Mental Number Line (MNL), with small numbers located on the left and larger ones on the right. How do these connections arise? Do we learn to associate numbers with space throughout cultural learning and social interactions or is this association rooted in the biology of the human brain? We showed that neonates spontaneously associate numbers with space. After being habituated to a certain number, neonates associated a smaller number with the left and a larger number with the right side. This evidence demonstrates that a predisposition to map numbers onto space is rooted in human neural systems.

---

Non-symbolic numerical skills are widespread in the animal kingdom (1). Pre-verbal infants (2,3) and non-human species (4) can extrapolate numerical magnitude from an array of elements, showing a non-symbolic number comprehension (5,6). In humans, this comprehension is present early in infancy (2) and can be assessed in adults whenever the use of language is prevented (7,8). Non-symbolic numerical tasks are easier, as the difference between the numbers increases (distance effect) and harder as the numerical magnitude increases (size effect), for both humans (9) and animals (8,10). These similarities are suggestive of a shared, ancient, non-verbal numerical mechanism (8). Therefore, uniquely human mathematical abilities seem to be based on a developmental and evolutionarily ancient “number sense” (11).

A peculiar characteristic of the numerical representation concerns the spatial coding of numbers along a left-right oriented continuum (12). Adults are faster at processing small numbers when responses are executed on the left side of space and faster for large numbers when responses are executed on the right side of space (spatial-numerical association of response codes, SNARC effect) (13). Several studies suggested that the left-to-right orientation of the mental number line is an out-come of exposure to formal instruction and that the mapping of number onto space would be a by-product of culture, based on reading/writing conventions and tool use, such as rulers (14). People for whom Arabic is the main language show an inverted SNARC effect (15), whereas people with mixed reading habits (i.e. those brought up reading both left-to-right and right-to-left) show no SNARC effect at all (16). However, an increasing number of studies support a phylogenetic origin of the mental number line. Seven-month-old infants looked longer at increasing (e.g. 1-2-3) but not at decreasing (e.g. 3-2-1) magnitudes displayed in a left-to-right spatial orientation (17). Eight-month-old infants oriented their attention toward the left after having seen a small number (i.e., 2), and toward the right after having seen a large number (i.e., 9) (18). Infant evidence excludes a primary influence of verbal counting in SNA orientation. However, this could be determined by the interactions with adults and the external world (19). A tendency to look longer at numerousness from left-to-right has been reported in our species (3,17). However, this is only a partial evidence of the SNA, because increasing the looking time from right-to-left has not been reported for decreasing sequences.

Adult Clark’s nutcrackers (20) and rhesus monkeys (21) have shown unilateral, left-to-right oriented bias to associate numerousness with space. Nevertheless, these biases could depend on continuous extents, which were not systematically controlled for. A complete evidence of a non-verbal SNARC-like phenomenon has been provided, up to now, only in domestic chicks, which preferentially respond to small numbers on the left side and to large numbers on the right side of space (22). Chicks associated a same non-symbolic number (i.e., an array of 8 squares) either with the left side, in the 8-32 range, or with the right side, in the 2-8 range. Such “relativity” of SNA is a fundamental characteristic of the human mental number line.

The underlying mechanism at the basis of chicks' SNARC-like effect might differ from the one that drives the effect in humans (16). Birds have laterally placed eyes, virtually complete nerve crossings at the optic chiasm and minimal interhemispheric connections, giving rise to a strong lateralization of function in everyday behavior (23). Humans, in contrast, like other primates, have frontally placed eyes, only partial crossing of nerves at the optic chiasm and strong interhemispheric connectivity. As a result, they show visual lateralization only in restricted conditions of vision (e.g. lateral presentation of briefly-presented stimuli) (24). The only way to discover the root of the human mental number line (20) is exploring whether human newborns, under minimal or no exposure to adults’ scanning biases, manifest SNA.

## Results

**Experiment 1: The origin of the SNARC effect.** Twenty-four full-term newborns, aged 51- hour-old (SD = 28.16, range: 12 – 117 h), took part at the Experiment 1 and were tested with an infant-control habituation technique.

Habituation and test stimuli consisted of static two-dimensional images. All stimuli contained a well-defined number of black square elements, average luminance 0.4 cd/m^2^, depicted on identical white square area of the dimension of 17.5 cm x 17.5 cm (694.68 pixels x 694.68 pixels), subtending a visual angle of 30.3° x 30.3° (average luminance 103 cd/m^2^). The number, the dimension and the position of the elements varied as a function of the experimental conditions. The distance elapsing between the closer edge of each stimulus and the center of the screen was of 4.25 cm (8.06°).

For the habituation phase we used five stimuli depicting 12 elements with a different spatial disposition. Each stimulus lasted 500 ms without any interval among stimuli presentation. We decided to employ five stimuli during the habituation phase in order to i) attract and maintain newborns’ attention and ii) prevent the newborns from learning to identify the stimuli on the basis of the spatial disposition of the black squares.

After the habituation phase with the number 12, a sequence of two different trials, counterbalanced between participants, was administered: half of the newborns had the small number in the first test trial and the large number in the second one (4-36), whereas the other half had the large number in the first test trial and the small number in the second one (36-4). In each test trial the same stimulus was simultaneously presented on the right and on the left side of a monitor. Each black square element measured 1.1 cm x 1.1 cm (43.67 pixels x 43.67 pixels), subtending a visual angle of 2.1° x 2.1°. Stimuli employed in the habituation phase were arrays composed of 12 black square elements. Test stimuli comprised a number of elements either smaller (4 black squares, for the small number test trial) or larger (36 black squares, for the large number test trial) than the number experienced during habituation (i.e., 12). In the present study, for the convenience of explanation, we calculated the percentage index using the stimulus on the left side for all experimental conditions. Therefore, scores significantly below 50% indicate a visual preference for the stimuli on the right side of the screen whereas, scores significantly above 50% indicate a preference for the stimulus on the left side.

All newborns included in the final sample reached the habituation criterion and they didn’t show any spatial biases, *t*_23_ = .56, *p* = 0.582 (M_left_ = 52.25%, see Fig.1).

**Figure 1.**
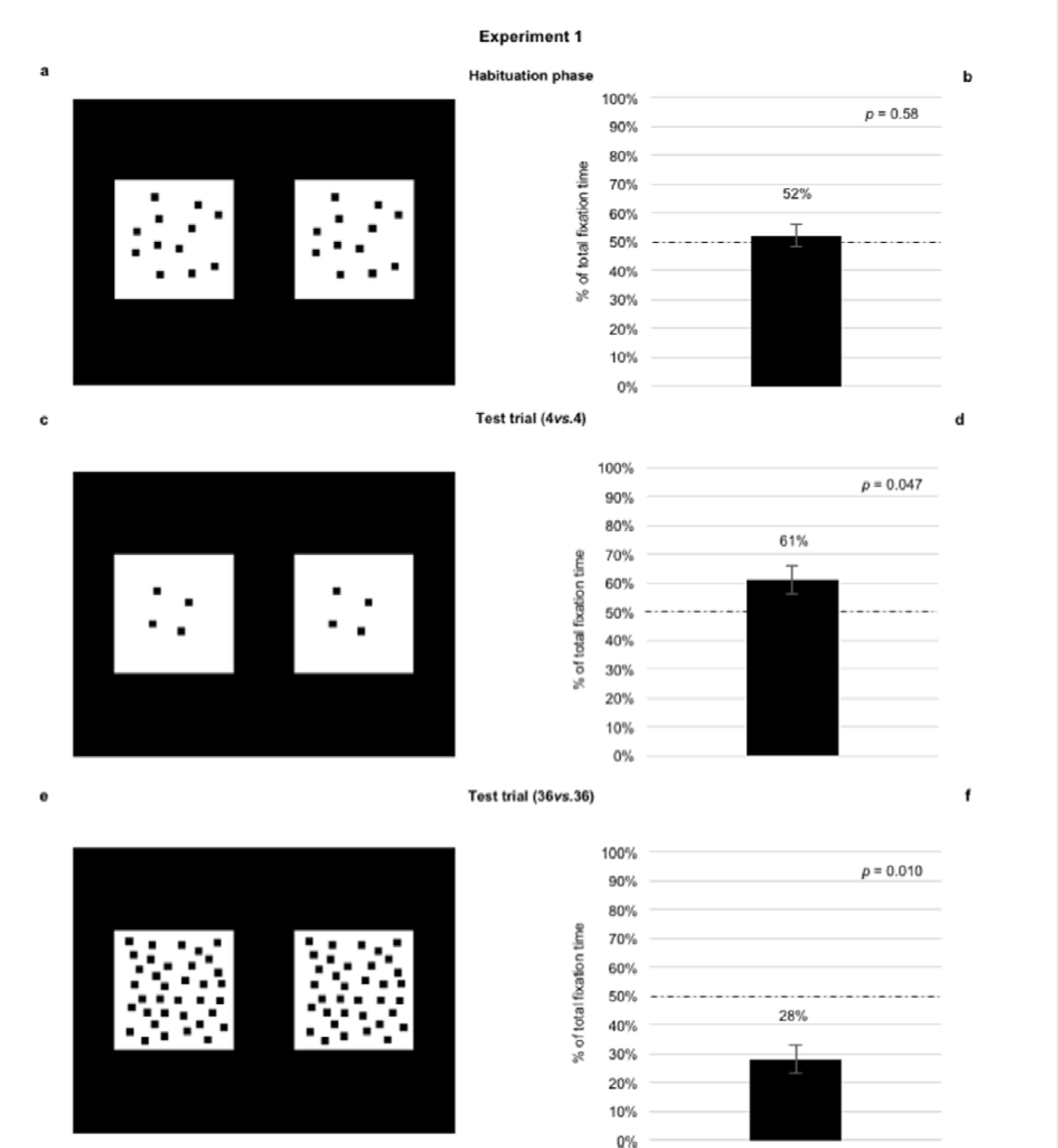
Stimuli and data on newborns’ visual preference 2 in Experiment 1. Newborns were habituated with two identical stimuli, depicting 12 black squares of the same dimension (a). Once they reached the habituation criterion, newborns underwent two test trials: one with a small number (4vs.4) (c), and the other with a large number (36vs.36) (e). During habituation, when the two stimuli depicted 12 squares, percentages of looking time toward the stimuli located on the left and on the right side of the screen did not significantly differ (b). In the test trial, when the two stimuli depicted 4 squares (4vs.4), newborns looked longer at the left-stimulus (d), when the stimuli depicted 36 squares (36vs.36), newborns looked longer at the right-stimulus (f). Errors bars are standard error and dashed lines indicate chance level (50%).

We carried out a repeated measure ANOVA with Test Trial Order (4-36 and 36-4) as a between-participants factor and Stimulus (4*vs*.4 and 36*vs*.36) as within-participants factor on the percentage of total fixation time toward left stimuli. The analysis revealed a significant main effect of Stimulus, *F*_1,22_ = 14.29, *p* < 0.001, 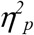 = 0.48 (number 4, M_left_ = 64.04%, SD = 18.65, *t*_23_ = 3.69, *p* = 0.001; number 36, M_left_ = 36.08%, SD = 22.85, *t*_23_ = −2.98, *p* = 0.007). Regardless of the Test Trial Order, newborns looked longer at the left-stimulus in the 4*vs*.4 trial, and at the right-stimulus in the 36*vs*.36 trial. However, since the first test trial could, theoretically, influence the second one, we analyzed only the first test trial. Results confirm that when the two stimuli depicted a number smaller than 12 (4*vs*.4, Fig.1), newborns looked longer at the left-stimulus (M_left_ = 61.25%, SD = 17.44, *t*_11_ = 2.24, *p* = 0.047), when the stimuli depicted a number larger than 12 (36*vs*.36, Fig.1), they looked longer at the right-stimulus (M_left_ = 28.00%, SD = 24.68, *t*_11_ = −3.09, *p* = 0.010).

These data suggest that at birth the association between small numerousness with the left side of space and large numerousness with the right side of space is already present. However, since the squares were identical in size, newborns’ preferences could have been driven by numerical or by continuous physical variables (overall perimeter and overall area).

**Experiment 2: The SNARC effect exhibits “relativity” in human newborns.** In Experiment 2a and 2b, we tested for two fundamental characteristics of the SNA in newborns: i) its independence from continuous physical variables; ii) its relative nature.

To exclude any possible use of continuous physical variables, we used squares of different dimensions during habituation and test trials. By controlling for the overall perimeter (the summation of perimeters of all squares depicted in both habituation and test stimuli was identical) we also controlled for the overall area (if the overall perimeter of two arrays of two-dimensional squares is identical, a negative correlation exists between numbers and overall area) (6).

To test SNA’s relativity, we habituated 12 neonates (Mean = 64.66 h, SD = 29.74, range 29 – 126 h; Experiment 2a) with the number 4 and a second group of 12 newborns (Mean = 52.75 h, SD = 42, range 11- 135 h, Experiment 2b) with the number 36. During a single test trial, both groups were presented with two identical stimuli, each depicting 12 squares, one on the left and one on the right side of the monitor.

In Experiment 2a and 2b, we equated the overall perimeter (i.e. summation of the perimeter of all black squares) and the overall area (i.e. summation of the area of all black squares) of the stimuli presented in the habituation and in the test trials. As a consequence, we obtained a negative correlation between the overall number of elements and their overall area.

As in Experiment 1, we used five stimuli with a different spatial disposition of the elements during the habituation phase. Specifically, in Experiment 2a, for the habituation phase we employed stimuli comprising of 4 black squares of 3.3 cm x 3.3 cm (131.50 pixels x 131.50 pixels), subtending a visual angle of 6.3° x 6.3°. The overall perimeter of the 4-elements was 58.2 cm. Test trial stimuli were 12 static black squares of 1.1 cm x 1.1 cm and therefore, with an overall perimeter of 58.2 cm. Importantly, the overall area of the 4-element stimuli (43.6 cm^2^) was larger than that of the 12-element stimuli (14.5 cm^2^). If the overall area, when the overall perimeter of the stimuli is identical, were the crucial factor underlying space-number association, newborns would have looked longer at the stimulus on the right side.

As in Experiment 2a, in Experiment 2b the overall perimeter between the habituation stimuli and the test stimuli was identical (158.4 cm). Habituation stimuli were 36 black squares (1.1 cm x 1.1 cm), whereas test stimuli were 12 static black squares, measuring 3.3 cm x 3.3 cm. The overall area of the 36-elements stimuli (43.6 cm^2^) was smaller than that of the 12-elements stimuli (130.7 cm^2^). If the overall area, when the overall perimeter of the stimuli is identical, were the crucial factor underlying space-number association, newborns would have looked longer at the stimulus on the left side.

All newborns included in the final sample reached the habituation criterion and they didn’t show any spatial biases either in Experiment 2a, *t*_11_ = .73, *p* = 0.480 (M_left_ = 54.25%, see Fig.2) and in Experiment 2b, *t*_11_ = .02, *p* = 0.980 (M_left_ = 50.17%, see Fig.3).

We ran a univariate ANOVA with Experiment (2a and 2b) as a between-participants factor on the percentage of total fixation time toward left stimuli. The analysis revealed a significant main effect of Experiment, F_1,22_ = 19.671, *p* = 0.001, 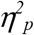 = 0.472. Neonates habituated with number 4 looked longer at the 12-elements on the right side (M_left_ = 30.17%, SD = 22.85, *t*_11_ = −3.01, *p* = 0.012, Fig. 2). Neonates habituated with number 36 looked longer at the 12-elements on the left side (M_left_ = 75.08%, SD = 26.62, *t*_11_= 3.26, *p* = 0.008, Fig. 3). These results show that SNA i) depends on number and not on other continuous physical variables, ii) is relative to the considered numerical range.

**Figure 2.**
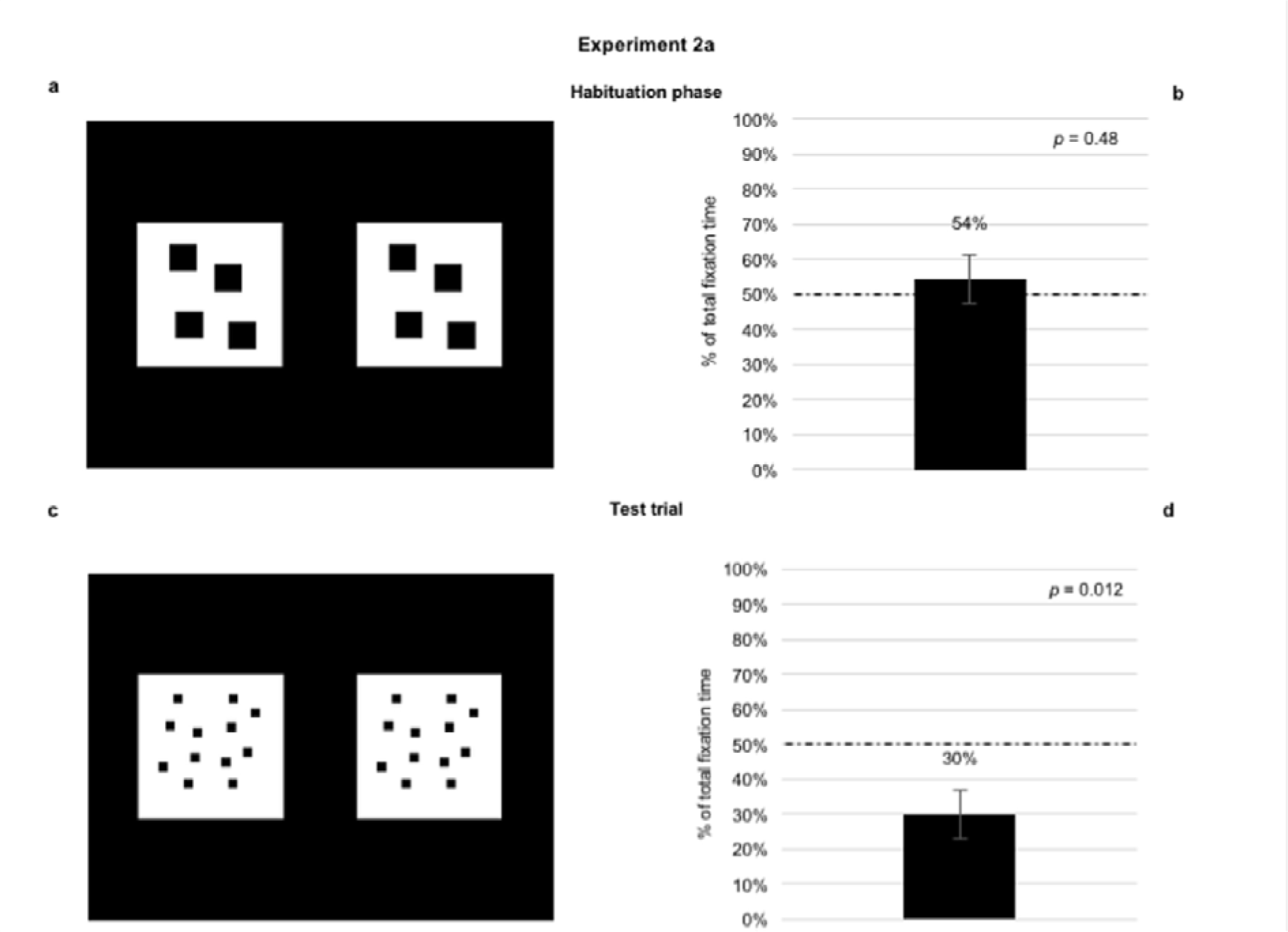
Stimuli and data on newborns’ visual preference 2 in Experiment 2a. In Experiment 2a we simultaneously controlled for the overall perimeter and area. We habituated a group of neonates with the number 4 (a) and then they were presented with the number 12 (12vs.12) in the test trial (c). As for the habituation phase, newborns did not show any visual preference (b). In the test trial, neonates looked longer at the right-stimulus (d). Error bars are standard error and dashed lines indicate chance level (50%).

**Figure 3.**
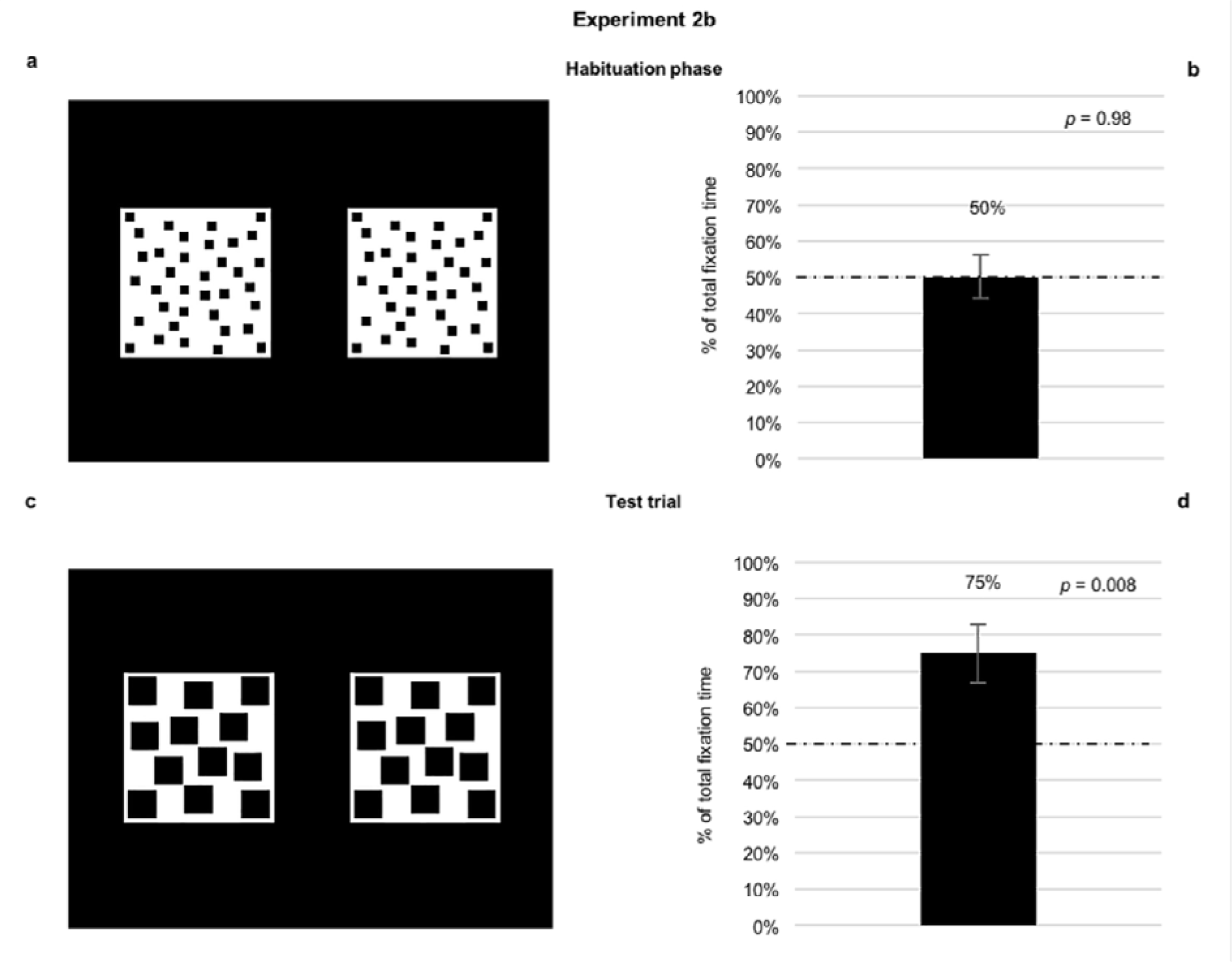
Stimuli and data on newborns’ visual preference in Experiment 2b. In Experiment 2b, as in Experiment 2a, we simultaneously controlled for the overall perimeter and area. We habituated a group of neonates with the number 36 (a) and then they were presented with the number 12 (12vs.12) in the test trial (c). As for the habituation phase, newborns did not show any visual preference (b). In the test trial, neonates looked longer at the left-stimulus (d). Error bars are standard error and dashed lines indicate chance level (50%).

## Discussion

The scientific dispute concerning the ultimate nature (cultural *vs*. biological) of the orientation of the mental number line is a strongly debated theoretical issue. On the one hand it has been suggested that it emerges as a result of exposure to formal instruction and culture (16,19). On the other hand, an increasing bulk of evidence has shown that pre-verbal infants and non-human animals associate numerousness with space, suggesting that the mental number line originates from pre-linguistic precursors (17, 18, 21, 37, 38, 39, 40). However, results obtained with infants could be accounted for either by innate or learning mechanisms. Up to now a complete association between small numbers and left space, and large numbers and right space has been provided solely in three-day-old domestic chicks (22). This evidence in completely inexperienced birds suggests that the role of reading and writing directionality is secondary in determining the orientation of the SNA (41, 21).

Caution has been urged, however, when using animal models to understand the origin of the orientation of human MNL (42). Convergent evolution, in which species from diverse evolutionary lineage could independently develop similar features (43), and strong differences in brain organization and lateralization (21, 44) could affect interpretation of comparative evidence (but see 45, 46). Nevertheless, comparative as well as developmental studies have been, so far, unable to unequivocally address the origin of human MNL. We overcame these limits by studying, for the very first time, human newborns with a very limited visual experience.

Here we provide evidence for a complete, relative and magnitude-based SNA in neonates. Hour-old newborns, initially habituated with a certain numerical value, spontaneously associated a smaller number with the left space and a larger number with the right space (Experiment 1). This association did not depend on the absolute magnitude of the number itself. Newborns habituated with number 4 associated the number 12 with the right (Experiment 2a), while newborns habituated with number 36 associated the number 12 with the left (Experiment 2b). This shows that SNA in newborns exhibits relativity.

Moreover, these findings cannot be explained by continuous physical variables. In fact, in Experiment 2a and in Experiment 2b, we controlled for the overall perimeter, obtaining an inverse correlation between overall area and number. Had newborns associated space to overall area, instead of number, their choices would have been the opposite to what reported.

The fact that day-old newborns rely on numerical rather than on quantitative information is in line with previous research which highlighted that, at the start of postnatal experience, we spontaneously use abstract numerical cues (3). Number is considered a fundamental perceptual feature that our brain process early to attain a complete representation of the external world (47, 48, 49, 50, 51, 52, 53; for a different perspective see 54). It seems that numerical competence did not emerge *de novo* in linguistic/symbolic adult humans, but was likely built on precursors available soon after birth (11, 55, 56). From this perspective, a non-symbolic number sense can be considered a developmental building block for the uniquely human capacity for mathematics (57). In support of this idea, it has been found that impairments to the non-symbolic numerical system are related to the occurrence of dyscalculia (58). The acuity of the non-symbolic numerical system is predictive of mathematical ability in early childhood (59) and throughout training it improves proficiency in symbolic mathematics (60). Our data strengthen the range of influence of non-symbolic numerical system on the symbolic one, showing that this affects also the directionality of the MNL.

Overall our findings show that SNA occurs with minimal experience, supporting the biological origin of SNA. This did not exclude that verbal (16) and non-verbal (19,25) experiences can modulate its directionality. Even if the orientation of the MNL reflects cultural effects (16), its widespread presence across diverse cultures supports the idea that the association between number and space is a universal cognitive strategy (61). Our evidence is strong, but also challenging. It is a starting point to disentangle the relative role and weight of cultural and biological factors in determining the orientation of the human mental number line.

## Acknowledgements

The authors are deeply indebted to Professors G. Perilongo, head of the Department of Health of woman and child and E. Baraldi, head of the Intensive Care and neonatal Pathology unit of the University of Padova. . We would also like to thank the nursing staff, the babies and their parents for their collaboration; Elena Berto, Daniela Carà and Silvia Dalò for assistance with new-born testing. A special thank to Kim Anne Barchi for the help during the editing of the manuscript.

This study was carried out within the framework of the agreement between the University and the Azienda Ospedaliera of Padova prot.: 91644645, and was supported by ‘Progetto di Ateneo’ 2012 to L.R. prot. CPDA127200) and by ERC Advanced Grant to G.V. (PREMESOR ERC-2011- ADG_20110406).

## Author Contributions

R.R., M.L., E.D.G., L.R. and F.S. designed the study; R.R., M.L. and E.D.G. created the stimuli; M.L. and E.D.G. collected, analyzed the data and did the statistical analyses; R.R., M.L., E.D.G., L.R., G.V. and F.S. interpreted the data, discussed the results; R.R., M.L. and E.D.G wrote the manuscript; L.R., G.V., B.D.B. and F.S. critically revised the manuscript.

## Author Information

The authors declare no competing financial interests.

Correspondence and requests for materials should be addressed to R.R.

## Methods

**Participants.** Sixty-nine (39 males) healthy, full-term, Caucasian newborns were selected from the maternity ward of the Pediatric Clinic of the University of Padova. Data from twenty-one babies (8 males) were discarded because i) they changed their state during testing (n = 10), ii) they showed a positional bias (n = 8), i.e. they looked in one direction for more than 80% of the time, and, iii) they were considered outliers (n = 3).

The final sample consisted of forty-eight (31 males) newborns randomly assigned to one of the three experimental conditions: twenty-four (10 males) took part in Experiment 1, twelve (6 males) in Experiment 2a and twelve (8 males) in Experiment 2b. All participants met the normal delivery screening criteria, had a birth weight between 2110 g and 4400 g (M = 3232.08 g, SD = 512.38), and an Apgar score of 9 at 5 min. Their mean postnatal age was 54.92 hours (SD = 32.26; range: from 17 to 135 hours). All newborns were tested only if awake and in an alert state^28^, and after the parents had provided informed consent. All experimental procedures were licensed by the Paediatric Clinic of the University of Padova (Protocol number 19147).

**Apparatus and procedures.** We conducted three experiments employing the infant-control habituation procedure^29,30^. Stimuli presentation and data collection were performed using E-Prime 2.0.

The baby sat on an experimenter’s lap and white curtains were drawn on both sides of the newborn to prevent interference from irrelevant distractors. The experimenter holding the baby was naive as to the hypothesis being tested and to the stimuli being presented and was instructed to fix her/his gaze on a monitor throughout the experimental session. Above the computer screen, the video camera recorded the eye movements of the newborns to control their looking behavior on-line and to allow off-line coding of their fixations.

At the beginning of each experiment, a red disc on a black background appeared to attract the newborn’s gaze to the center of the monitor. In a continuous fashion, the disc changed in size from small (1.8 cm) to large (2.5 cm) until the newborn’s gaze was properly aligned with the red disc. The red disc blinked at a rate of 300 ms on and 300 ms off. The sequence of trials was then started by a second experimenter who watched the newborn’s eyes through the monitor. When the newborn’s gaze was aligned with the red disk, the second experimenter pressed a keyboard key that automatically turned off the central disc and activated the onset of the stimuli, thereby initiating the sequence of trials.

The paradigm comprises the habituation phase and the test trials, in which two identical stimuli were presented side-by-side. In the habituation phase, stimuli remained on the screen until the habituation criterion was reached. Newborns were judged to have habituated when, from the fourth fixation onward, the sum of any three consecutive fixations was 50% or less of the total of the first three fixations^31^. A bilateral, rather than a central, presentation was selected for two reasons: i) when newborns look at a centrally presented stimulus, it is difficult for a coder to decide whether they are actually looking at the stimulus or simply not moving their eyes from the central position; ii) at birth, the photoreceptors in the central fovea are very immature, resulting in poor vision in the central area of the visual field^32,33^.

During the test trials, one of the experimenters video recorded the newborns’ visual behavior on each stimulus. A test trial ended when newborns did not fixate on the display for at least 10 s.

The newborns sat on an experimenter’s lap at a distance of 30 cm from the computer screen. The stimuli were displayed on an Apple LED Cinema Display (Flat Panel 30") computer monitor (refresh rate = 60 Hz, resolution 2560 x 1600 pixels).

**Data Scoring and Statistical Analyses.** Two coders, unaware of the stimuli presented, analyzed off-line, the videos by coding newborn’s eye movements frame by-frame.

The dependent variables that we measured, as in previous studies^34,35^, were some of those suggested by Cohen’s model of attention^36^ as indexes of the attention-getting and of the attention-holding mechanisms. As indexes of attention-getting mechanisms, the coders recorded separately for each stimulus and each position, the direction of the first fixation and the number of orientations for the two stimuli. This latter measure was then converted to a percentage score by computing the total number of orientations towards a given stimulus divided by the total number of orientations towards both stimuli in each visual preference condition, X 100. As indexes of attention-holding mechanisms, the coders recorded the longest fixation and the total fixation time (i.e., the sum of all fixations) towards the two stimuli. Also in the case of the total fixation time, we calculated the percentage of visual preference, that is, the length of time for which each newborn looked at a given stimulus divided by the total time spent looking at both stimuli in each test trial, X 100.

The mean estimated reliability between observers on the 64.58% of the overall participants (i.e., 31 out of 48 newborns) was Pearson’s r = 0.93, p < 0.001.

Data of Experiment 1 were analyzed as follow:

i) a first analysis was conducted on the percentage of total fixation time towards the left position during the habituation phase, to control for any spontaneous a-priori preference for a specific position (i.e., spatial biases);

i) a repeated measure ANOVA with Test Trials Order of stimuli presentation (4-36 and 36-4) as a between-participants factor and Stimulus (4*vs*.4 *versus* 36*vs*.36) as within-participants factor on the percentage of total fixation time and number of orientations toward left stimuli was carried out;
ii) a two-sided one sample t-test on percentage of the number of orientations and a binomial test on the direction of the first fixation in the first test trial;
iii) a two-sided one sample t-test on percentage of total fixation time and a two-sided paired t-test (left *vs*. right) on the duration of the longest fixation in the first test trial.

In Experiment 2a and 2b, we run the following analyses:

i) we replicated the same analysis carried out in Experiment 1 during the habituation phase to control for any spontaneous a-priori preference for a specific position (i.e., spatial biases);

i) a univariate ANOVA with Experiment (2a and 2b) as a between-participants factor on the percentage of total fixation time and number of orientations toward left stimuli was carried out, in order to test relativity of SNA;
ii) a two-sided one sample t-test on percentage of the number of orientations and a binomial test on the direction of the first fixation were carried out;
iii) a two-sided one sample t-test on percentage of total fixation time and a two-sided paired t-test (left *vs*. right) on the duration of the longest fixation, were carried out.

Data were analyzed using SPSS software.

For a complete description of results, see SI.

### Data availability

The datasets are available from the corresponding author on reasonable request.

## References

1. Vallortigara G (2014) Foundations of number and space representations in non-human species. In Evolutionary Origins and Early Development of Number Processing, eds Geary D-C, Bearch D-B & Mann Koepke K (Elsevier, New York), pp 35–66.

2. Cordes S, Brannon E-M (2009) Crossing the divide: Infants discriminate small from large numerosities. Dev Psychol 45(6):1583–1594.

3. Izard V, Sann C, Spelke E-S, Streri A (2009) Newborn infants perceive abstract numbers. Proc. Natl Acad Sci USA 106 (25):10382–10385.

4. Vallortigara G (2012) Core knowledge of object, number, and geometry: a comparative and neural approach. Cogn Neuropsychol 29 (1-2):213–236.

5. Feigenson L, Dehaene S, Spelke E-S (2004) Core systems of number. Trends Cogn Sci 8(7):307–314.

6. Rugani R, Castiello U, Priftis K, Spoto A, Sartori L (2017) What is a number? The interplay between number and continuous magnitudes. Behav Brain Res in press.

7. Cordes S, Gelman R, Gallistel C-R, Whalen J (2001) Variability signatures distinguish verbal from nonverbal counting for both large and small numbers. Psychon Bull Rev 8(4):698–707.

8. Cantlon J-F, Brannon E-M (2007) Basic Math in Monkeys and College Students. PLoS Biol 5, e328.

9. Moyer R-S, Landaeur T-K (1967) Time required for judgments of numerical inequality. Nature 215:1519–1520.

10. Scarf D, Hayne H, Colombo M (2011) Pigeons on Par with Primates in Numerical Competence. Science 334:1664.

11. Dehaene S. (2011) The number sense: How the mind creates mathematics, revised and updated edition (New York, Oxford University Press).

12. Galton F (1880) Visualised numerals. Nature 21:252–256.

13. Dehaene S, Bossini S, Giraux P (1993) The mental representation of parity and number magnitude. J Exp Psychol Gen 122:371–396.

14. Rugani R, de Hevia M-D (2017) Number-space associations without language: Evidence from preverbal human infants and non-human animal species. Psychon Bull Rev 24:352–369.

15. Zebian S. (2005) Linkages between number concepts, spatial thinking and directionality of writing: The SNARC effect and the REVERSE SNARC effect in English and in Arabic monoliterates, biliterates and illiterate Arabic speakers. J Cog Cult 5:165–190.

16. Shaki S, Fischer M-H, Petrusic W-M (2009) Reading habits for both words and numbers contribute to the SNARC effect. Psychon Bull Rev 16:328–331.

17. de Hevia M-D, Izard V, Coubart A, Spelke E-S, Streri A (2014) Representations of space, time, and number in neonates. Proc Natl Acad Sci USA 111(13): 4809–4813.

18. Bulf H, de Hevia M-D, Macchi-Cassia V (2015) Small on the left, large on the right: Numbers orient preverbal infants’ visual attention onto space. Dev Sci 19(3):394–401.

19. Patro K, Fischer U, Nuerk H-K, Cress U (2016) How to rapidly construct a spatial–numerical representation in preliterate children (at least temporarily). Dev Sci 19(1):126–144.

20. Rugani R, Kelly D-M, Szelest I, Regolin L, Vallortigara G (2010) Is it only humans that count from left to right? Biol Lett 6(3):290–292.

21. Drucker C-B, Brannon E-M (2014) Rhesus monkeys *(Macaca mulatta)* map number onto space. Cognition 132:57–67.

22. Rugani R, Vallortigara G, Priftis K, Regolin L (2015) Number-space mapping in the newborn chick resembles humans’ mental number line. Science 347(6221):534–536.

23. Vallortigara G, Versace E (2017) Laterality at the neural, cognitive, and behavioral levels. In APA Handbook of Comparative Psychology: Vol. 1. Basic Concepts, Methods, Neural Substrate, and Behavior, eds Call J (American Psychological Association, Washington DC), pp 557–577.

24. Ocklenburg S (2017) Tachistoscopic Viewing and Dichotic Listening. In Lateralized Brain Functions, eds Rogers L-J, Vallortigara G (Springer Verlag, New York), pp 3–28.

25. Bächtold D, Baumüller M, Brugger P. (1998) Stimulus–response compatibility in representational space. Neuropsychologia 36:731–735.

26. Harvey B-M, Klein B-P, Petridou N, Dumoulin S-O (2013) Topographic Representation of Numerosity in the Human Parietal Cortex. Science 341(6150): 1123–1126.

27. Ditz H-M, Nieder A (2015) Neurons selective to the number of visual items in the corvid songbird endbrain. Proc Natl Acad Sci USA 112(25):7827–7832.

28. Prechtl H, O’Brien M-J (1982) Behavioral states of the full term newborn: The emergence of a concept. In Psychobiology of the human newborn eds Stratton P. (New York, Wiley), pp 53–73

29. Horowitz F-D, Paden L, Bhama K, Self P (1972) An infant-control procedure for studying infant visual fixation. Dev Psychol 7(90).

30. Sokolov E-N (1963) Perception and the condition reflex (New York, Macmillan).

31. Slater A, Earle D-C, Morison V, Rose D (1985) Pattern preferences at birth and their interaction with habituation-induced novelty preferences. J Exp Child Psychol 39:37–54.

32. Abramov I, et al (1982) The retina of the newborn human infant. Science 217:265–267.

33. Atkinson J, Braddick O (1989) Development of basic visual functions. In Infant development eds Slater A, Bremner G (London, Erlbaum), pp 7–41.

34. Valenza E, Simion F, Macchi-Cassia V, Umiltà C (1996) Face preference at birth. J Exp Psychol Human 22:892–903.

35. Di Giorgio E, Leo I, Pascalis O, Simion F (2012) Is the face-perception system human-specific at birth? Dev Psychol 48:1083–1090.

36. Cohen L-B (1972) Attention-getting and attention-holding processes of infant visual preferences. Child Dev 43:869–879.

37. Lourenco S-F, Longo M-R (2010) General magnitude representation in human infants. Psychol Sci 21(6):873–881.

38. Adachi I (2014). Spontaneous spatial mapping of learned sequence in chimpanzees: Evidence for a SNARC-like effect. PLoS One 9: e90373.

39. Rugani R, Rosa-Salva O, Regolin L (2014) Lateralized mechanisms for encoding of object. Behavioral evidence from an animal model: the domestic chick (*Gallus gallus*). Front Psychol 5.

40. Rugani R, Vallortigara G, Regolin L (2016) Mapping number to space in the two hemispheres of the avian brain. Neurobiol Learn Mem 133:13–18.

41. Brugger P (2015) Chicks with a number sense. Science, 347(6221): 477–478.

42. Patro K, Fischer U, Nuerk H-K, Cress U (2016) How to rapidly construct a spatial–numerical representation in preliterate children (at least temporarily). Dev Sci 19(1): 126–144.

43. Emery N-J, Clayton N-S (2004) The mentality of crows: convergent evolution of intelligence in corvids and apes. Science, 306(5703):1903–1907.

44. Fischer M-H, Shaki S (2016) Measuring spatial–numerical associations: evidence for a purely conceptual link. Psychol Res 80(1): 109–112.

45. Rugani R, Vallortigara G, Priftis K Regolin L (2015) Response to Comments on ”Number-space mapping in the newborn chick resembles humans’ mental number line”. Science 348:1438.

46. Rugani R, Vallortigara G, Priftis K Regolin L (2016) Response: “Newborn chicks need no number tricks. Commentary: Number-space mapping in the newborn chickresembles humans' mental number line”. Front Hum Neurosci 10:31.

47. Burr D, Ross J (2008) A visual sense of number. Curr Biol 18(6):425–428.

48. DeWind N-K, Adams G-K, Platt M-L, Brannon E-M (2015) Modeling the approximate number system to quantify the contribution of visual stimulus features. Cognition, 142:247–265.

49. Anobile G, Cicchini G-M, Burr DC (2016) Number as a primary perceptual attribute: a review. Perception 45 (1–2), 5–31.

50. Cicchini G-M, Anobile G, Burr D-C (2016) Spontaneous perception of numerosity in humans. Nature Commun, 7: 12536.

51. Fornaciai M, Cicchini G-M, Burr D-C (2016) Adaptation to number operates on perceived rather than physical numerosity. Cognition 151: 63–67.

52. Fornaciai M, Brannon E-M, Woldorff M-G, Park J (2017) Numerosity processing in early visual cortex. NeuroImage 157:429–438.

53. Park J, DeWind N-K, Woldorff M-G, Brannon E-M, (2016) Rapid and direct encoding of numerosity in the visual stream. Cereb Cortex 26: 748–763.

54. Leibovich T, Ansari D (2016) The symbol-grounding problem in numerical cognition: A review of theory, evidence, and outstanding questions. Can J Exp Psychol 70(1):12.

55. Carey S (2009) The origin of concepts (Oxford University Press).

56. Vallortigara G (2017) An animal’s sense of number. In The nature and development of mathematics. Cross disciplinary perspective on cognition, learning and culture, eds Adams J-W, Barmby P, Mesoudi A (Routledge, New York), pp 43–65.

57. Spelke E-S (2000). Core knowledge. Am Psychol 55: 1233–1243.

58. Wilson A-J, Dehaene S (2007) Number sense and developmental dyscalculia. Hum Behav Learn Dev Brain: Atypical Dev 2: 212–237.

59. Starr A, Libertus M-E, Brannon E-M (2013) Infants show ratio-dependent number discrimination regardless of set size. Infancy 18: 927–941.

60. Park J, Brannon E-M (2013) Training the approximate number system improves math proficiency. Psychol Sci 24: 2013–2019.

61. Göbel, S-M, Shaki S, Fischer M-H (2011) The cultural number line: A review of cultural and linguistic influences on the development of number processing. J Cross-Cultural Psychol 42: 543–565.

